# Extrapolating acute bee sensitivity to insecticides using a phylogenetically informed interspecies scaling framework

**DOI:** 10.1101/2020.05.05.078204

**Authors:** Tobias Pamminger

## Abstract

Plant protection products, including insecticides, are important for global food production. Historically, research of the adverse effects of insecticides on bees has focused on the honeybee (*Apis mellifera*), while non-*Apis* bee species remained understudied. Consequently, sensitivity assessment of insecticides for the majority of bees is lacking, which in turn hinders accurate risk characterization and consequently bee protection. Interspecies sensitivity extrapolation based on body weight offers a potential solution to this problem, but in the past such approaches have often ignored the phylogenetic background and consequently non independence of species used in such models. Using published data on the sensitivity of different bee species to commonly used insecticides, their body weight and phylogenetic background I build interspecies scaling models (ISMs) applying a phylogenetically informed framework. In addition, I compared, the relative sensitivity of the standard test species *Apis mellifera* to other bee species to evaluate their protectiveness when used as standards screening bee species in the risk assessment process. I found that overall 1) body weight is a predictor of bee sensitivity to insecticides for a range of insecticide classes and 2) *A*. *mellifera* is the most sensitive standard test species currently available and consequently a suitable surrogate species for ecotoxicological risk assessment.

## Introduction

Insects, in particular bees, play a vital role in pollinating wild plants as well as crops [1]. This ecosystem service provided by both domesticated (e. g. *A. mellifera*) and wild bees, is essential for maintaining the functionality of non-managed (e g. biodiversity), as well as food production in agricultural landscapes [2, 3]. Due to their ecological and economical importance, reports of wild bee population declines in some areas have raised concerns regarding the sustainability of pollination services [4–6]. Several stressors including changes in land use, intensified agricultural practice and bee diseases are being discussed as drivers for this dynamic [7–9]. Within this broad discussion, the use of PPPs, in particular insecticides, and their potential adverse side effects on wild bee populations has emerged as a controversially discussed topic [9].

In order to accurately estimate the potential risk for bees following insecticide application in an agricultural setting, a robust risk assessment is needed [10]. A prerequisite for such a process is accurate hazard (sensitivity) characterization (e g. lethal doses or rates) of insecticides for the bees in question, which together with realistic exposure estimates, provides the basis for a robust estimation of the actual risk bees face. However, besides a few well-studied model bee species (e g. *A. mellifera* and *Bombus terrestris*) little is known about the response of the majority of bees to insecticides to date. As *A. mellifera* is currently used as the main surrogate species for non-Apis bees risk assessment around the globe, this lack of comparative sensitivity data generates uncertainty regarding the protectiveness of this approach [11–13].

The problem of limited toxicological information and the need for inter-species extrapolation, in the context of risk assessment, is not new and different approaches have been developed to address it in other areas including toxicology [14–16]. A straightforward method is cross species sensitivity extrapolation based on body weight, which is current practice in other areas (e g. vertebrates) of PPP risk assessment (e.g. US EPA wildlife risk assessment [17]). In vertebrates, it is assumed and generally accepted that toxicity, similarly to other fundamental traits (e g. metabolic rate), is closely linked to the size and consequently weight of an organism with smaller ones being more susceptible [16, 18]. As many scaling relationships seems to apply to most living organisms [18–20] it is very likely that toxicity scaling extends beyond vertebrates and can be applied to bees as well.

Besides size, other factors likely play a role in shaping the sensitivity of an organism to a given substance. Such factors include extrinsic ones such as the type and properties of the substance (e g. water solubility) and intrinsic properties like receptor composition affecting molecule binding strength [21] and effectiveness of detoxification pathways [16, 21, 22]). Such intrinsic factors are likely linked to the organisms’ evolutionary history (phylogeny) as more closely related species are often more similar in their traits compared to more distantly related ones [23]. This important and potentially confounding factor for such inter-species analyses has long been recognized [24], but only recently have authors started to directly address this issue in their ecotoxicological studies [25–28].

In this paper I use a phylogenetically informed analytical framework to build inter-species scaling models (ISMs) using acute contact toxicity for the currently most commonly used insecticide classes and bee body weight to investigate, 1) whether scaling relationships can be used to extrapolate bee sensitivity based on their body weight, 2) if scaling relationships differ between different classes and 3) to which degree scaling relationships are shaped by the underlying phylogenetic relationship among species. In a last step 4), I investigate the relative sensitivity of bees of the Apis genus compared to other bees and discuss its consequences for surrogate bee species selection in the context of ecotoxicological risk assessment.

## Materials and methods

### Data collection

Between June 2017 and July 2019 I conducted a literature search used Google Scholar and Scopus (last search June 2019) and the search terms bee* AND tox* AND LD50. I screened the returned publications and their respective reference section for relevant publications. A paper was considered relevant if it met the following criteria: 1) reporting acute contact toxicity data (LD_50_/bee) following direct exposure paradigm similar to the OECD 214 guidelines [29] (2) testing an insecticides (active substance or formulation) and 3) using a bee species (Apiformes) as test species. In case all criteria where met the results were included in the analysis.

### Data selection and aggregation

All reported substance specific LD_50_ were categorized according to their mode of action (MoA [30]) and the four currently most commonly used insecticides were selected for further analysis [31–34]. The neonicotinoids were split between cyano- (c-) and nitro- (n-) substituted neonicotinoids, as it is known that bee species’ responses can drastically differ between these two classes [22, 35, 36]. This resulted in five insecticide classes for analysis: acetylcholinesterase inhibitors (AChE-inhibitors), pyrethroids, c- and n-substituted neonicotinoids and organochlorines. After an initial screen one paper Del Sarto et al. [37] was excluded from further analysis because the results reported for *A. mellifera* deviated drastically (up to 700 times higher LD_50_) from all other results reports for the same substance-species pair.

For each publication in the remaining dataset I calculated the geometric mean per chemical class resulting in one endpoint per data source (paper) and chemical class. As quality and study design drastically varies between publications this approach was chosen to limit the disproportionate influence of individual studies on the overall results without excluding data. For the same reason I calculated the geometric mean as the arithmetic mean is highly susceptible to outliers [38]. In a last step, I calculated the geometric mean per bee species for all insecticide classes which are the values used in the final analysis. For all bee species present in the final dataset I collected data on bee wet weight in milligram as a predictor variable for bee sensitivity. In cases where I did not find species-specific weight data or multiple reported values for a species, I used the average reported weight for the genus or the species respectively. The final dataset thus contained bee species, insecticide class, bee weight and LD_50_ values. Because I assumed a power law or power law like association between the variables commonly encountered in body mass associated traits including sensitivity to xenobiotics [18–20], all used acute contact LD_50_ values and bee wet weights were log_10_ transformed prior to analysis.

### Phylogeny

For all bee species represented in the dataset I constructed an initial phylogeny using the “rotl” R package [39]. “rotl” uses data from the open tree of life project (https://tree.opentreeoflife.org) to reconstruct the phylogenetic relationship of the input species. Using this phylogeny as a back bone I used recent publications on bee phylogeny [40–44] to resolve some remaining polytomies. Because there are no consistent branch length information available for the resulting tree I Grafen-transformed [45] the tree using the R package “ape” [46].

### Phylogenetic signal

In a first step I tested for phylogenetic signal in both bee weight and LD_50_ of the five insecticide classes separately. Phylogenetic signal is used as an indicator to see if more closely related species are more similar in the respective trait than expected by chance. I used the phylosig() function in the R package “phytools” using the “K” estimation method [47].

### Interspecies scaling models (ISM)

A linear model (LM) was fit for every insecticide class independently with LD_50_ values as the response, bee wet weights as the predictor and number of publications as a weighting factor in order to account for the uneven sampling coverage between species (see Tab S1). In order to assess the robustness of the model outputs (e g. slope) the same models were 1) fitted without the weighting factor and 2) with the weighting factor but excluding *A. mellifera*. Function fitting was performed using ordinary least square regression implemented in the *lm* function in R. Residuals were inspected visually for normality (QQ plot), homogeneity of variances (residuals versus predicted plot).

### Phylogentic ISM diagnostic

Following the procedures outlined by [48], I checked the residuals of each final linear model for phylogenetic signals using the phylosig() function in the “phytools” package [47] using the “K” method. In cases where a significant phylogenetic signal is detected it is suggested [48] to use a phylogenetically controlled general least square regression (PGLS) as implemented in e g. the “caper” package [49].

### Weight corrected LD_50_ & relative sensitivity

In order to visualize and compare the LD_50_ across different insecticide classes and bee species I calculated the weight-corrected LD_50_ (LD_50_/g bee wet weight) and tested if they exhibited phylogenetic signal and compared the results to the primary analysis of the corresponding uncorrected LD_50_ values. In order to assess the overall sensitivity of a given species across insecticide classes, I calculated the standardized (z-transformed) LD_50_/g bee for all bee species, from now on referred to as relative sensitivity, for all five insecticide classes. This approach is necessary because different subset of bees have been sampled for the insecticide classes (Tab. S1) and if uncontrolled for one might simply compare the median size of bees tested per insecticide class instead of the actual toxicity (See Fig. S1 A & B). For all species where I had information on more than three insecticide classes, I additionally calculated their overall median standardized relative sensitivity.

### Graphics

All analyses were conducted in R v 3.5.1 [50]. Graphs and trait mapping on the phylogeny were done using ggplot2 [51] and phytools [47], respectively.

## Results

In total, I collected data from 55 data sources containing 343 reports of acute toxicity LD_50_ measurements. The final data set provided information for 28 bee species belonging to 12 genera spanning five of the seven currently recognized bee families [41]. However, the data was heavily skewed towards AChE-inhibitors (N = 164) and the model species *Apis mellifera* (N = 131). For a summary of the results and associated references see Tab. S1 & 2, ESM 1 & 2).

### Phylogenetic signal

I found clear evidence for phylogenetic signal for bee weight (K = 0.71, p = 0.001) and the LD_50_ for AChE-inhibitors (K = 0.3, p = 0.016) suggesting that more closely related species are more similar for these two traits. In contrast I did not find evidence for a phylogenetic signal for any of the other insecticide class specific LD_50_ measurements (all K < 0.6, all p > 0.32).

### Interspecies scaling models

In four out of five insecticide classes, I found evidence that bee LD_50_ are positively associated with body weight (all R^2^ > 0.27 see Fig. 1A-C & E) meaning that heavier bees have higher LD_50_ (are less sensitive). This relationship was statistically significant for three of the classes (AChE-inhibitors, c-neonicotinoids and pyrethroids) and a trend for n-neonicotinoids (p = 0.09). For organochlorines I found no clear indication for an association (R^2^ = 0.12, p = 0.36 Fig. 1D). When looking at the estimated regression slopes, I find that c-neonicotinoids likely scale allometrically with body weight (the 95% CI of slope did not overlap with 1) while the slope for all other insecticide classes was consistent with isometric scaling (95% CI of slope overlap with 1). The slope estimates were robust when removing *A. mellifera* or using an unweighted model (see Tab. S3). Except for organochlorines the 95% CI did not overlap with 0 further supporting an overall positive association between bodyweight and LD_50_ (see Tab. S3).

**Figure 1:**
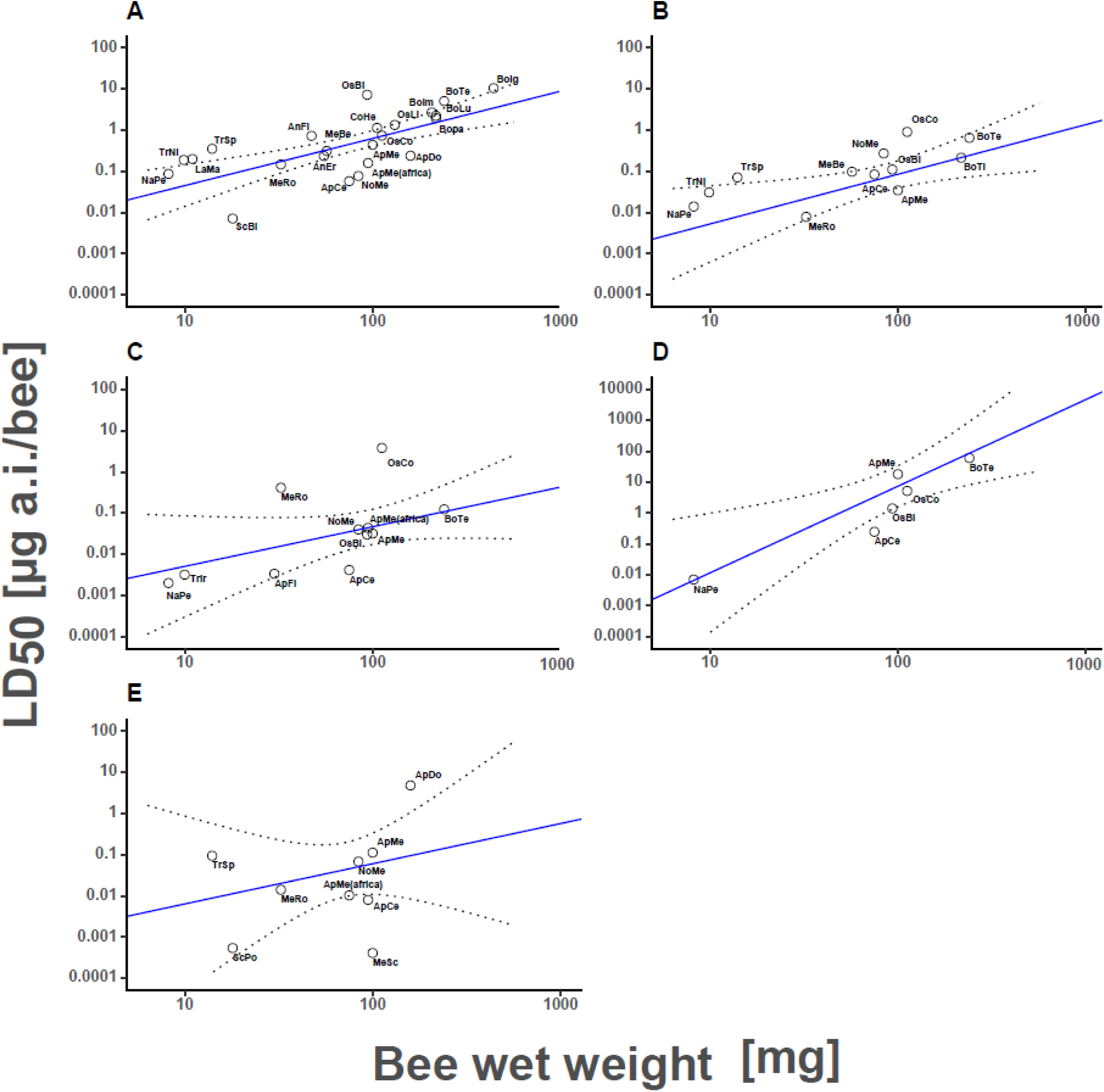
Depicts the association between the log_10_ transformed bee body weight and sensitivity for the five insecticide classes (LM regression line and 95%CI). We show this association for AChE inhibitors (A), pyrethroids (B), n-neonicotinoids (C), c-neonicotinoids (D) and organochlorines (E). The numerical results for the LM can be found in Table 1.

### Phylogenetically controlled interspecies scaling models

I found no indication for phylogenetic signal in the model residuals (all p > 0.32 indicating that fitting a linear model without correcting for phylogenetic autocorrelation can be considered robust in this case [48] which rendered a phylogenetically controlled approach (e g. PGLS) unnecessary.

**Table 1:**
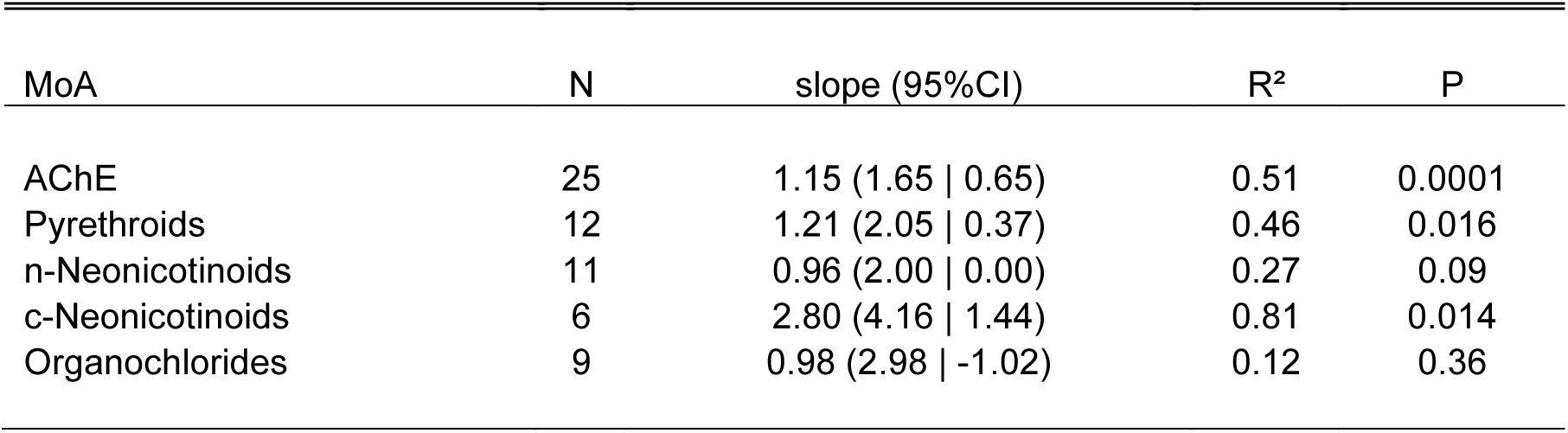
Summary results of the LM fit to the log_10_ bee weight and sensitivity. We present samplesize (number of bee species), slope and 95% CI (Upper bound | Lower bound), R^2^ and associated P values

### Weight corrected LD_50_ & relative sensitivity

When visualizing the weight-corrected LD_50_ (log_10_ LD_50_ / g bee) values of bee species across the different chemical groups, I found that at least some species of the *Apis* genus are always among the most sensitive (Fig. 2 A-E). While in some cases non-*Apis* bee species are more sensitive compared to Apis bees e g. *Scaptotrigona bipunctata* (see Fig 2A), *Megachile rotundata* (see Fig. 2B) and *Melipona scutellaris* (see Fig. 2E) it has to be noted that these values are based on single studies making these estimates less reliable (see Fig.S2). A notable exception to this overall picture were c-neonicotinoids (Fig. 2D) where *A. mellifera* was among the most resilient species together with *B. terrestris*.

**Figure 2:**
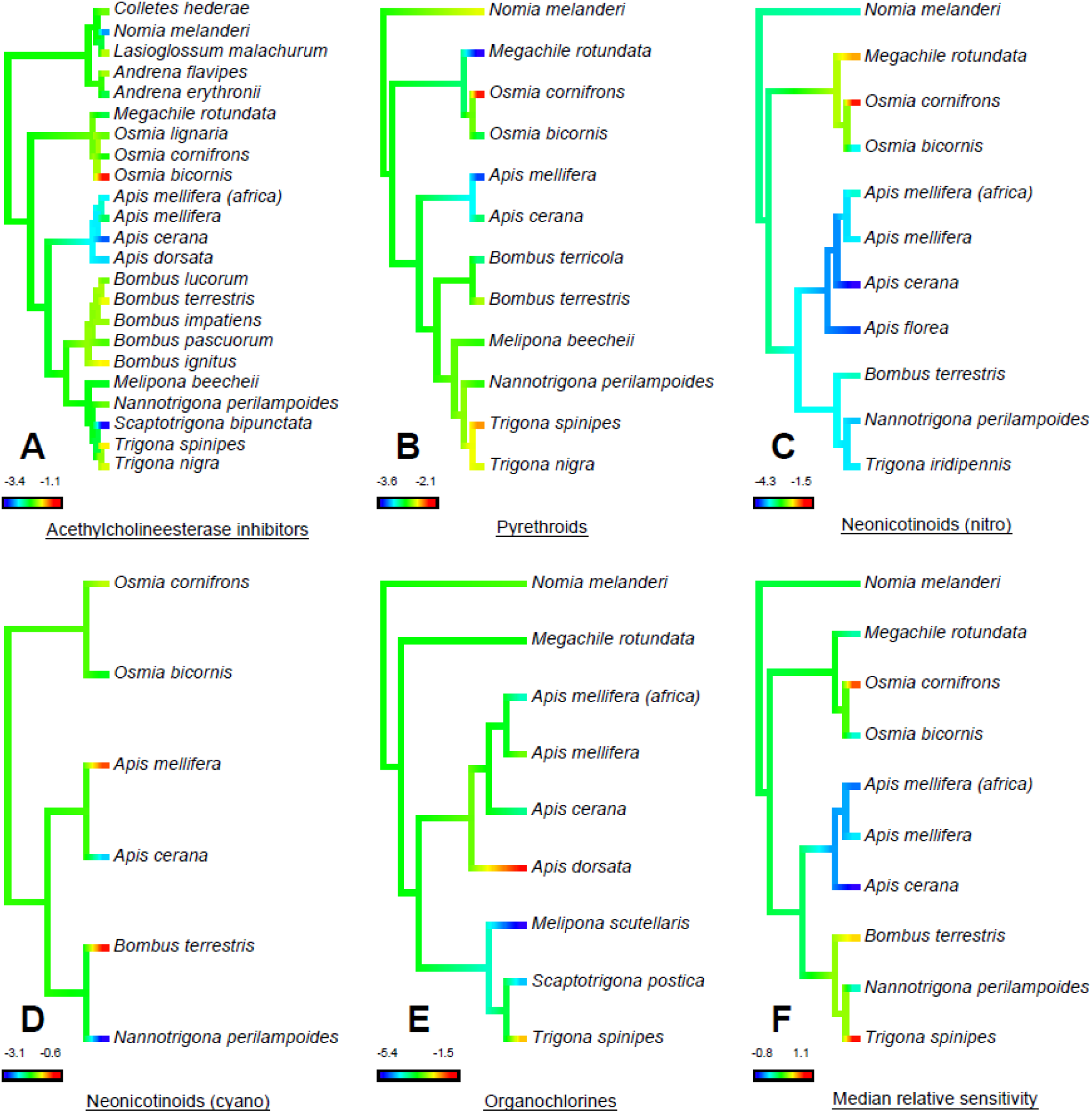
Depicts the log_10_ transformed LD_50_ per gram bee for the five insecticide classes plotted on the bee phylogeny: AChE inhibitors (A), pyrethroids (B), n-neonicotinoids (C), c-neonicotinoids (D) and organochlorines (E). In the last penal the median relative sensitivity (z-transformed LD_50_ per gram bee) for all species where data on more than 2 insecticide classes was available (all relative sensitivity values can be found in Table S4).

When looking at the relative sensitivity, I found that overall bees of the *Apis* genus were the most sensitive (Fig. 2F, Tab.2) when considering all insecticide classes. In contrast bees of the genus *Bombus, Osmia and Trigona* rank among the more resilient bees found in this analysis (Fig.2, Tab.2 & Tab. S4). When looking at the neotropical stingless bees we find a diverse picture *Scaptotrigona* being more sensitive compared to other bees such as *Trigona*. However, these results are based on a very limited number of studies (Fig. S2E). For none of the insecticide classes I found significant phylogenetic signal for the weight corrected LD_50_ (All p > 0.34).

**Table 2:** Standardized relative sensitivty of bee species where informtaion for more than three MoA

| Scientific Name | Relative sensitivity | N |
| --- | --- | --- |
| *Apis_cerana* | -0.83 | 5 |
| *Apis_mellifera* | -0.31 | 5 |
| *Apis_mellifera_(africa)* | -0.54 | 3 |
| *Bombus_terrestris* | 0.69 | 4 |
| *Megachile_rotundata* | -0.13 | 4 |
| *Nannotrigona_perilampoides* | -0.21 | 4 |
| *Nomia_melanderi* | 0.10 | 4 |
| *Osmia_bicornis* | -0.23 | 4 |
| *Osmia_cornifrons* | 0.95 | 4 |
| *Trigona_spinipes* | 1.07 | 3 |

## Discussion

The available data indicates that body weight, similarly to vertebrates [15, 16], is an important and often strong predictor of acute bee sensitivity towards insecticides. However, the strength of the scaling relationship (slope) can differ between insecticide classes. When taking body weight into account, *Apis mellifera* bees seem particularly susceptible to a range of common insecticide classes (with the notable exception of c-neonicotinoids for *A. mellifera*), while *Osmia* and *Bombus* bees appear comparatively resilient.

### Weight corrected LD_50_ Relative sensitivity & Phylogenetic patterns

When plotting the weight corrected LD_50_ measurements and the relative sensitivity on the phylogeny an interesting pattern emerged. In the majority of cases members of the Apis genus are among the most sensitive species present in the data set (see Fig. 2). The idea that *Apis* bees are particularly sensitive has been present in the literature for some time with some evidence supporting it [11–13, 52, 53]. It is tempting to speculate about the underlying cause for this phenomenon. One might argue that domestication and the strong anthropogenic driven selection for certain traits (e.g. increased honey yields and low aggression) might have unintentionally resulted in increased sensitivity to xenobiotics. However, the fact that not only the domesticated honeybee *A.mellifera*, but all members of the genus including feral ones (*e g. Apis florea*) share this increased susceptibility hints towards a different explanation. In contrast to most other bees all species of the *Apis* genus are highly eusocial [54–56]. This means that these species live in groups and reproduction is monopolized by a single female (queen) with all other members of the colony being non-reproductive helpers (workers; see [55]). Currently standard ecotoxicological toxicity testing is done using the worker class and several studies have shown that reproductive status strongly affects the bees’ behavior and physiology including sensitivity to some toxic substances [57]. For example, *A.mellifera* workers have been reported to be up to 54 (mean 15) times more sensitive to some xenobiotics than queens [58]. While worker sterility might be part of the explanation of the elevated sensitivity compared to many solitary and therefore reproductively active bees (e g. *Osmia bicornis*), it might not fully explain why workers other semi- and eusocial worker bees (e g. *Trigona spinipe* and *B. terrestris*) appear resilient to a range of insecticide classes (Fig. 2). However, in both case worker bees are known to reproduce and lay unfertilized eggs which develop into males [59, 60] in particular once the queen is absent [61] similarly to the standard test conditions in the laboratory [62, 63]. If reproductive activity provides some protection from adverse effects of insecticides (e g. improved detoxification) this might explain the observed elevated resilience. It might be a promising approach to investigate and compare the reproductive status of workers of other eusocial bee species in order to better understand the observed patterns of sensitivity.

In addition to this physiological explanation there is some genomic evidence suggesting that *A. mellifera*, and by extension likely additional members of the *Apis* genus, lack some substantial detoxifying enzymes (including glutathione‐S‐transferases, cytochrome P450 monooxygenases and carboxyl/cholinesterases) which are essential for detoxifying a range of xenobiotics including insecticides [53]. This line of evidence is supported by recent findings which pinpoint members of the P450 family as key determinates of bee sensitivity to different insecticides [22, 35, 36]. When taken together these results support the previous suggestion that *A. mellifera* is sensitive [11] and consequently a sensible option [12] to use as standard surrogate test species for contact scenarios during the risk assessment process.

### Scaling patterns and scaling coefficients

Our results suggest that it is in principal possible to predict the acute sensitivity of bee species to insecticides based on their body weight. While this holds true for four out of five chemical classes of insecticides investigated, the organochlorines are a potential exception. It is possible that organochlorine acute toxicity in bees doesn’t scale with body weight at all, but given that many scaling relationships hold across a wider taxonomic range [16, 19, 20, 64], it is more likely that this relationship is obscured by possible outliers likely caused by experimental conditions which is a substantial problem when overall sampling size is low. In order to resolve this issue, additional species that are not included in the current data set could be tested and/or outlier species have to be re-tested by independent labs.

However, when looking at the slope associated large confidence intervals it becomes clear that there is still a substantial amount of uncertainty associated with the estimated scaling coefficients (Tab.1 & Tab. S4) likely due to low sampling size and variation in data quality. Before such approaches can be considered for actual risk assessment purposes more work is needed to reduce the uncertainties associated with these models and test if its predictions hold for bee species not present in the current data set. In order to effectively improve the model predictions, it would be fruitful to focus future exploratory bee toxicity testing on 1) species-insecticide class combinations not present in the data set and 2) focus primarily on bee species occupying the edges of weight the parameter space (very small and very large bees) in order to limit the need of model extrapolation outside its parameter space.

### Risk assessment considerations

Accurate sensitivity (hazard) characterization is at the heart of current ecotoxicological risk assessment [65]. In the case of bees, the honeybee has traditionally been used as a surrogate test species for this purpose. However, the relative sensitivity of honeybees compared to other bee species is not well characterized and consequently it is unclear if it is a suitable surrogate system for the approximately 20 00 bee species worldwide [54].

There have been two approaches to improve overall sensitivity characterization for non *Apis* bees: 1. Develop new model test species (e g. *Bombus terrestris* and *Osmia bicornis*) for lower tier screening or 2) better characterize the relative sensitivity of *A. mellifera* in relation to other bees and develop and appropriately conservative risk assessment based on the honeybee [11–13].

Our study suggests that the second approach would be feasible as *Apis* (including *A. mellifera*) worker bees, compared to other bees, appear to be highly sensitive to a range of different insecticide classes in acute contact scenarios. In combination with the extensive experience and resulting robust test methodology makes *A. mellifera* solid choice as screening level test species. In contrast the relative sensitivity data indicates that the two recently developed model species *Bombus terrestris* and *Osmia bicornis* for the European risk assessment process are comparatively resilient to a range of insecticides, which in turn means that including them in the screening step will likely not make the risk assessment more protective for bees from a sensitivity perspective [12, 13, 66, 67].

One could imagine a scenario, once robust scaling relationships have been established, to use scaling coeffects to extrapolate the sensitivity of an insecticide class to other bees based on their bodyweight and the respective honeybee sensitivity. This approach would circumvent the problem that it is unlikely to ever test enough bee species to adequately cover non Apis bee sensitivity (~20 000 species globally) and limit excessive bee testing necessary to cover all regulatory requirements.

### Merging ecotoxicological and evolutionary thinking

When conducting comparative analyses, it is important to consider the non-independence of the different species due to their phylogenetic associations [24]. As Revell [44] pointed out, the often used two-step-approach: 1. testing for a phylogenetic signal in the response or predictor variable and in case of a significant signal apply 2. phylogenetically controlled approaches (PIC, PGLS etc.) is misguided and does not address the underlying technical problem of correlated model residuals potentially violating model assumptions [48]. Revell rightly suggested to only control for phylogeny if the model residuals show phylogenetical signal violating LM assumptions. However, in case of limited sampling this approach will likely suffer from low power to detect phylogenetic signal. In such cases an evolutionary perspective becomes important when deciding the analytical approach. In a first step it should always be considered if we expect phylogenetic signal to occur. One important aspect in this regard is the scale of the analysis. When comparing more distantly related taxa it becomes more likely to find more pronounced differences in a trait of question and consequently phylogenetic signal. In the case of insecticides, we can observe such pattern for n-neonicotinoids. When comparing mammals and insects (phyla) we find clear sensitivity differences with mammals being less susceptible compared to insects [21]. This effect is likely driven by compound specific variation in the binding strength to the receptor [21]. However, when looking within insects (order) or even at lower taxonomic levels (e g. Apiformes) we find less pronounced absolute differences. In such a case it would make sense to use phylogenetically controlled approaches in the former, but less so in the latter case.

In the case of this analysis I found that both bee sensitivity (LD_50_) to AChE-inhibitors and bee weight exhibit phylogentic signal while the weight corrected LD_50_ values do not. This suggests that the phylogenetic signal found for AChE-inhibitors sensitivity is likely driven by bee body weight. This finding, in combination with the lack of phylogenetic signal in the model’s residuals, supports the use of a simple LM for the analysis [48]. However, it also shows the importance of using an evolutionary framework when deciding on the most appropriate analysis for a given data set. Without such considerations one might be tempted to simply fit LMs, even when LM assumptions are violated, which could facilitate the misinterpretation of such data [48].

In recent years an increasing number of comparative statistical tools as well as phylogenetic information became available to researchers (e g. “rotl”, “phytools” and “ape” [39, 46, 47]). It is now technically feasible to conduct such analyses while addressing the potential pitfalls associated with inter-species analysis. When carefully thinking about the analytical approach these tools will increase the robustness and reliability of inter-species inference, which is particularly important and exciting for larger data sets. In the context of ecotoxicology the awareness for the need of a phylogenetic informed analytical framework has been recognized [25–27, 68] and the tools available will facilitate the further integration of evolutionary thinking in ecotoxicology.

### Conclusions and future directions

1. When taking body weight into account, we find across most insecticide classes, genus *Apis* (including *A. mellifera*) seem particularly sensitive to a range of insecticides during contact exposure.
2. This pattern (see 1), emerging on a genus level, the clear association of bee weight and phylogeny and the association between AChE enhibitors phylogeny all highlights the importance of taking the evolutionary background of species into account when interpreting inter-species results.
3. Our results show that similarly to other biological systems, body weight is an important predictor of bee acute contact sensitivity to a range of insecticide classes and could be used to extrapolate bee sensitivity to non-tested species.
4. In order to improve the sensitivity prediction based on bodyweight it is important to focus future exploratory bee sensitivity testing afford. This afford should be focused on 1) so far untested species-insecticide class pairs 2) independent confirmation of extrema results (“potential outliers”) and 3) on species on end of the weight spectrum (heavy and light) to minimize the necessity to extrapolate outside the parameter space.
5. In the future, I hope that toxicity data for other exposure pathways (oral, chronic and developmental) will become available which in turn will provide a more complete picture of bee sensitivity across the phylogeny and ultimately could improve the reliability of bee risk assessment.

## Acknowledgements

I would like to thank Matthias Bergtold, Christof Schneider and Magdalena Mair and the terrestrial ecotoxicology team at BASF for in-depth discussions on the topic and careful reading of previous versions of the manuscript.

---

**Figure S1:**
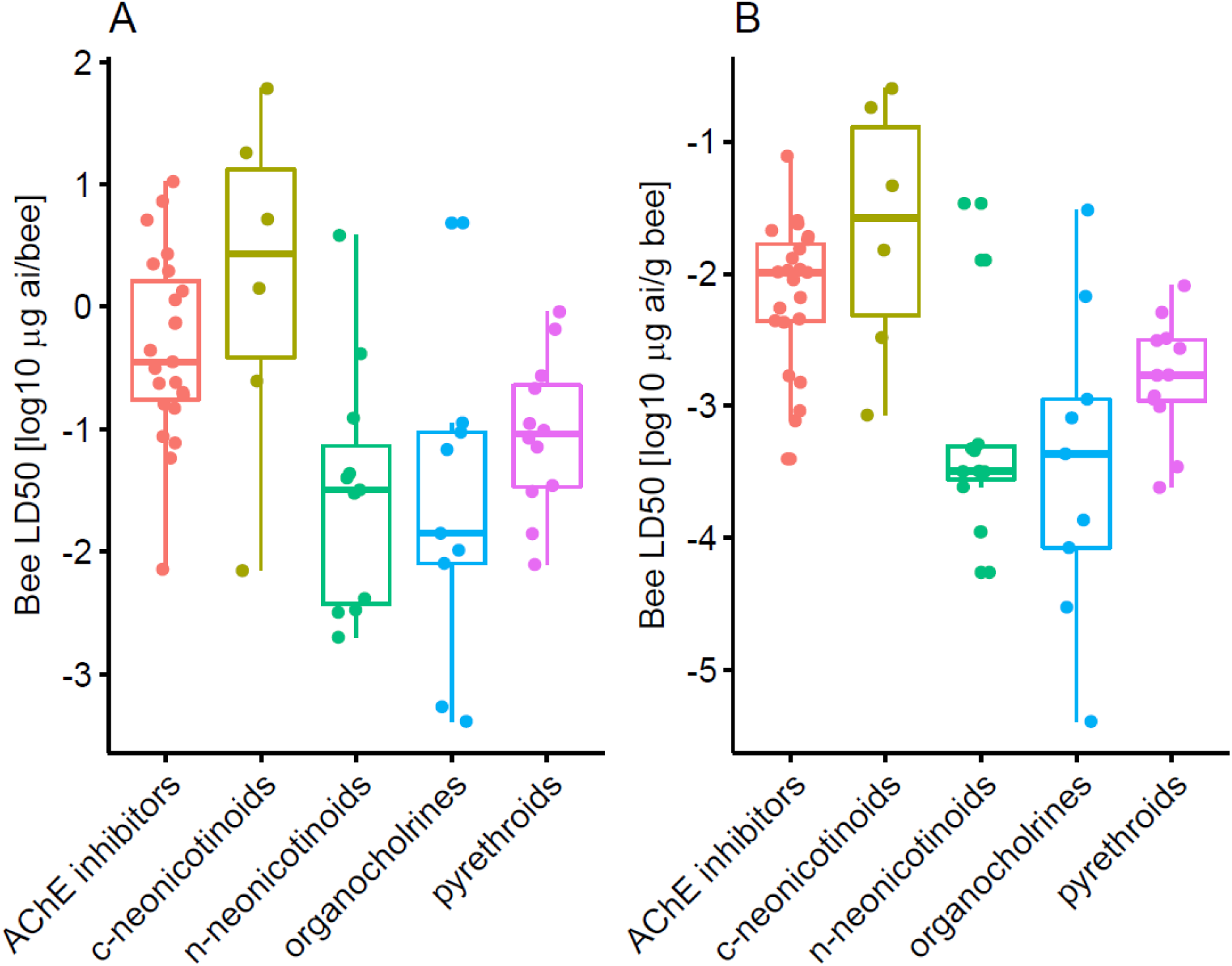
Comparison of uncorrected (A) and weight corrected (B) sensitivity of bees to exposure of five different classes of insecticides.

**Figure S2:**
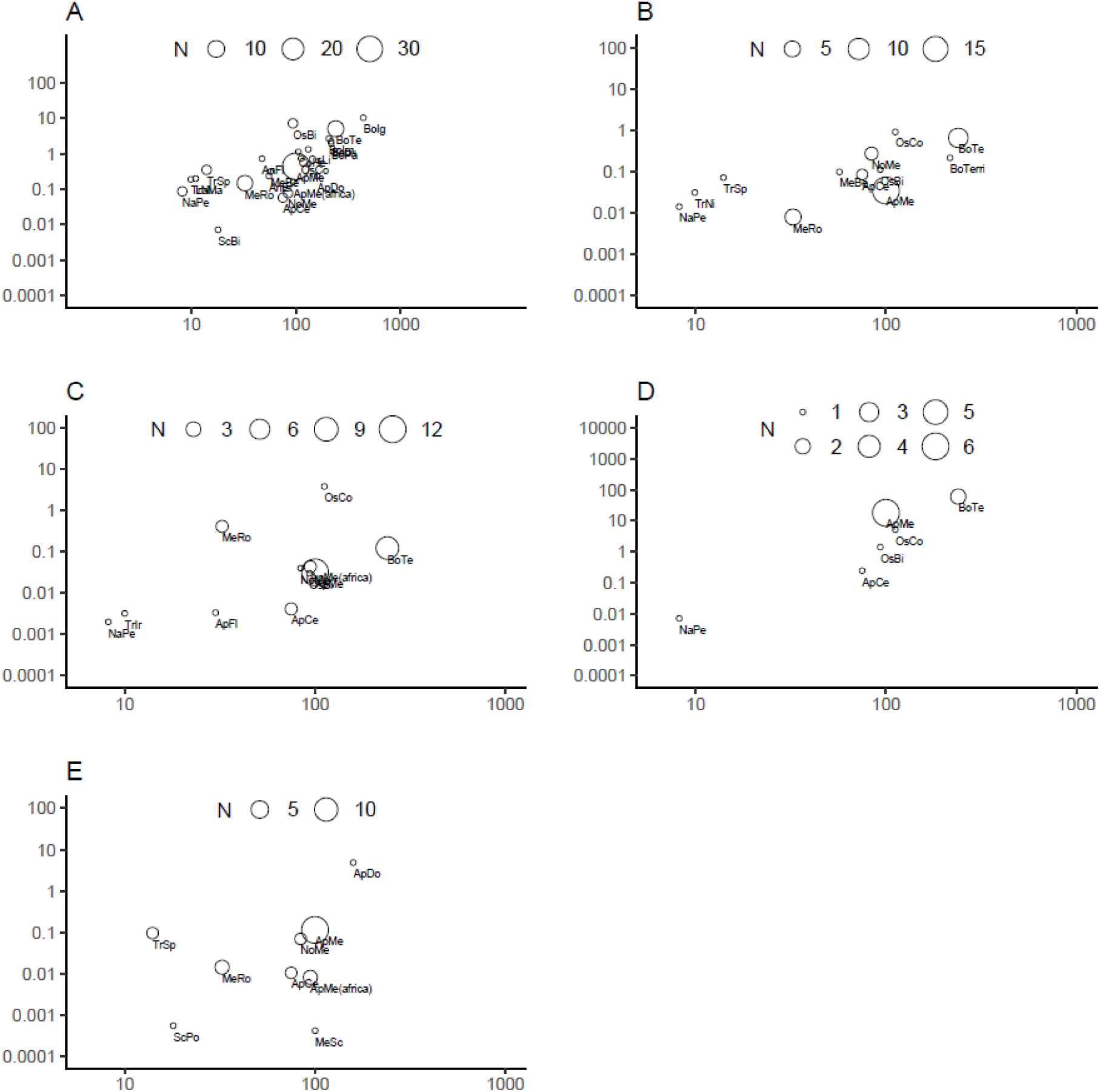
Depicts the association between the log_10_ transformed bee body weight and sensitivity for the five insecticide classes (similar to Fig. 1) We show this association for AChE inhibitors (A), pyrethroids (B), n-neonicotinoids (C), c-neonicotinoids (D) and organochlorines (E). Circle sizes indicate the number of bees species samples for this data point.

**Table S1:**
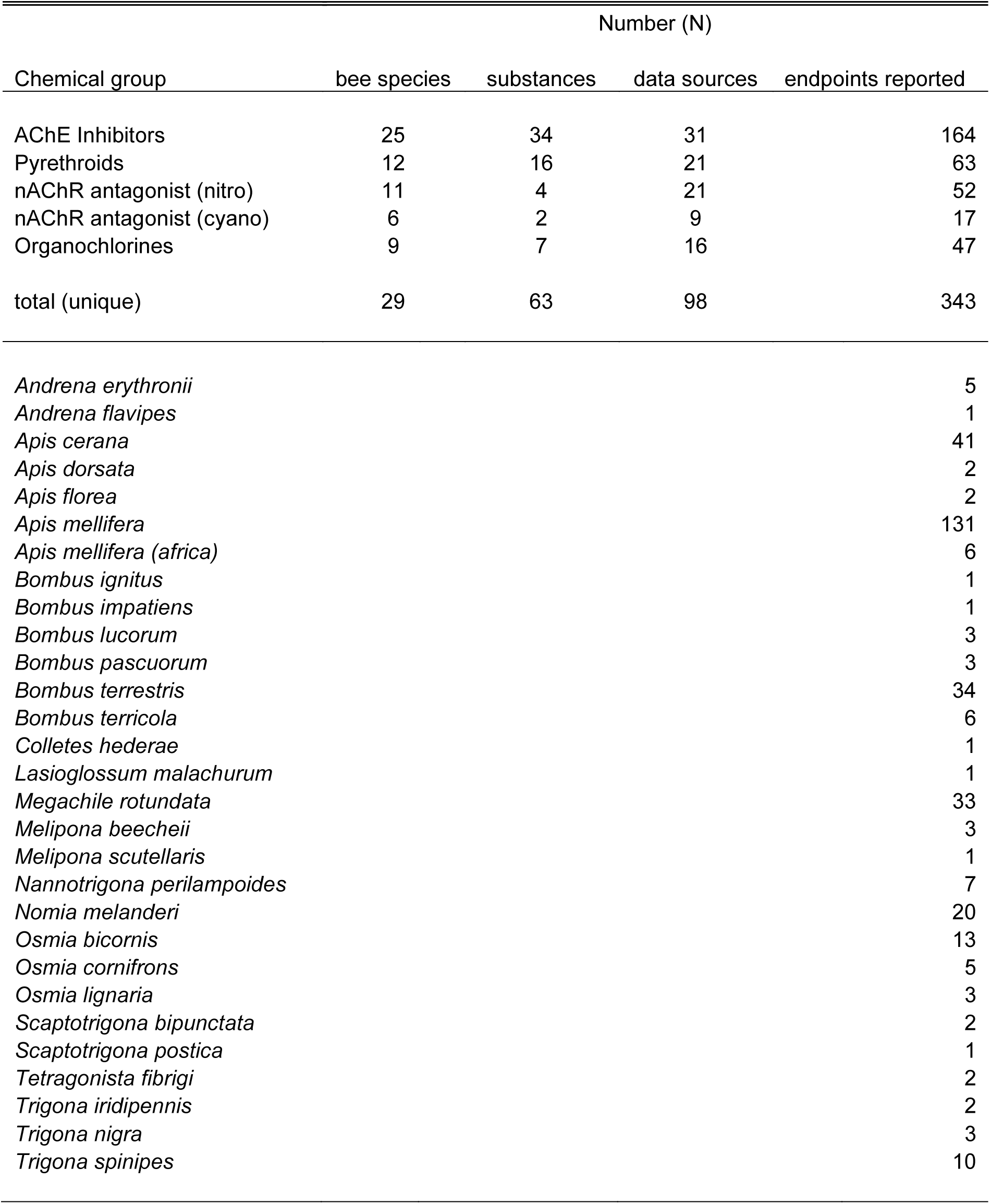
Summary statistics of the collected dataset regarding number bee species, substances and results found.

**Table S2:**
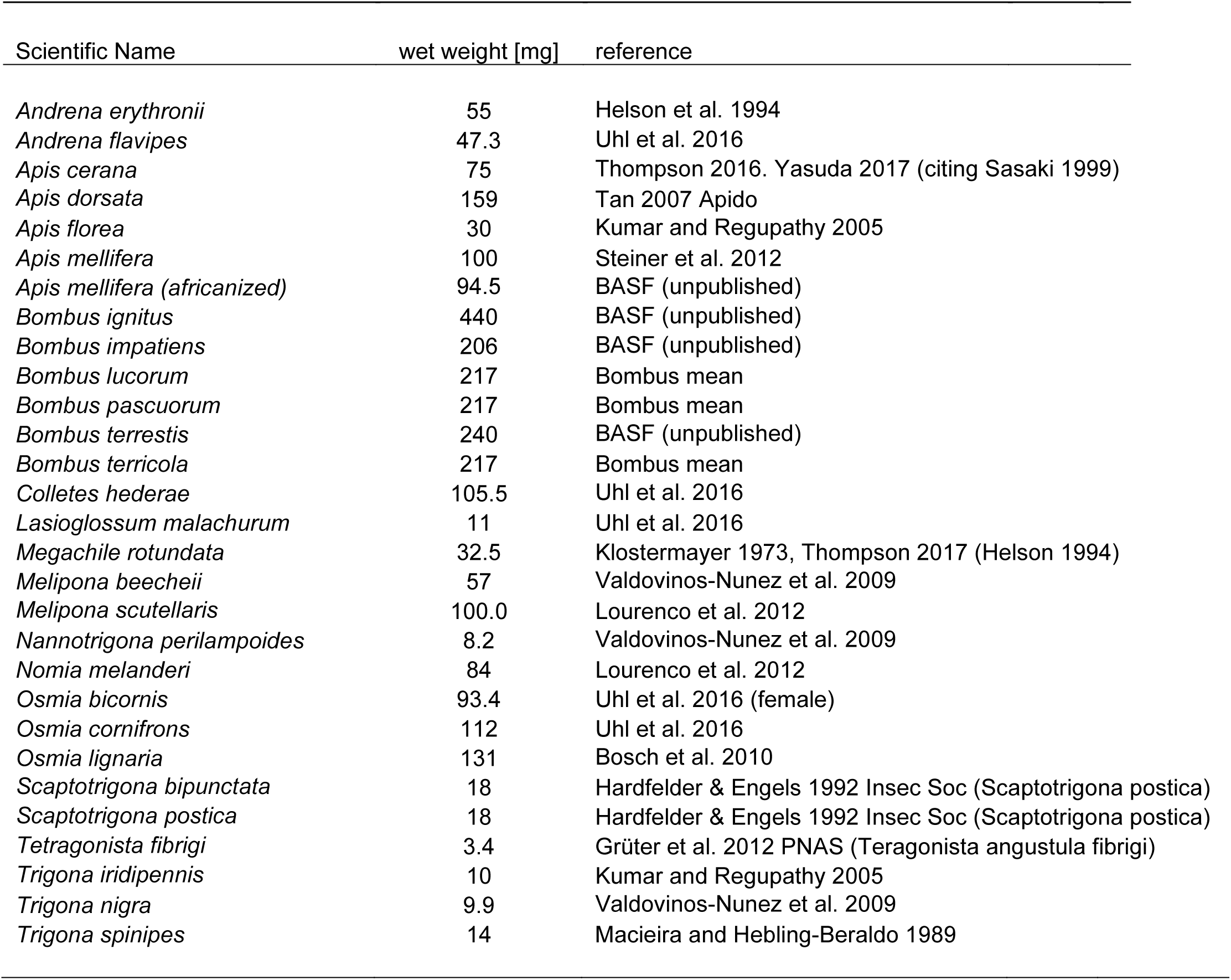
Wet weight of bees in mg and associated references the full list is found in ESM 2.

| Scientific Name | wet weight [mg] | reference |
| --- | --- | --- |
| *Andrena erythronii* | 55 | Helson et al. 1994 |
| *Andrena flavipes* | 47.3 | Uhl et al. 2016 |
| *Apis cerana* | 75 | Thompson 2016. Yasuda 2017 (citing Sasaki 1999) |
| *Apis dorsata* | 159 | Tan 2007 Apido |
| *Apis florea* | 30 | Kumar and Regupathy 2005 |
| *Apis mellifera* | 100 | Steiner et al. 2012 |
| *Apis mellifera (africanized)* | 94.5 | BASF (unpublished) |
| *Bombus ignitus* | 440 | BASF (unpublished) |
| *Bombus impatiens* | 206 | BASF (unpublished) |
| *Bombus lucorum* | 217 | Bombus mean |
| *Bombus pascuorum* | 217 | Bombus mean |
| *Bombus terrestris* | 240 | BASF (unpublished) |
| *Bombus terricola* | 217 | Bombus mean |
| *Colletes hederæ* | 105.5 | Uhl et al. 2016 |
| *Lasioglossum malachurum* | 11 | Uhl et al. 2016 |
| *Megachile rotundata* | 32.5 | Klostermayer 1973, Thompson 2017 (Helson 1994) |
| *Melipona beecheii* | 57 | Valdovinos-Nunez et al. 2009 |
| *Melipona scutellaris* | 100.0 | Lourenco et al. 2012 |
| *Nannotrigona perilampoides* | 8.2 | Valdovinos-Nunez et al. 2009 |
| *Nomia melanderi* | 84 | Lourenco et al. 2012 |
| *Osmia bicornis* | 93.4 | Uhl et al. 2016 (female) |
| *Osmia cornifrons* | 112 | Uhl et al. 2016 |
| *Osmia lignaria* | 131 | Bosch et al. 2010 |
| *Scaptotrigona bipunctata* | 18 | Hardfelder & Engels 1992 Insec Soc (Scaptotrigona postica) |
| *Scaptotrigona postica* | 18 | Hardfelder & Engels 1992 Insec Soc (Scaptotrigona postica) |
| *Tetragonista fibrigi* | 3.4 | Grüter et al. 2012 PNAS (Teragonista angustula fibrigi) |
| *Trigona iridipennis* | 10 | Kumar and Regupathy 2005 |
| *Trigona nigra* | 9.9 | Valdovinos-Nunez et al. 2009 |
| *Trigona spinipes* | 14 | Macieira and Hebling-Beraldo 1989 |

**Table S3:** Assessment of the robustness of the slope estimates for each mode of action for the implemented linear models (“main model”: including weights = number of publications all data). I compared it to 1) the estimates of a model without weights (“excluding weights”) and 2) to a model including weights after the removal of *Apis mellifera* (“excluding *A. mellifera”*).

| MoA | main model | excluding weights | excluding *A. mellifera* |
| --- | --- | --- | --- |
| AChE | 1.15 (1.65 \| 0.65) | 1.04 (1.54 \| 0.54) | 1.23 (1.73 \| 0.73) |
| Pyrethroids | 1.21 (2.05 \| 0.37) | 0.89 (1.45 \| 0.33) | 1.35 (1.97 \| 0.73) |
| n-Neonicotinoids | 0.96 (2 \| 0) | 1.28 (2.48 \| 0.08) | 0.98 (2.06 \| -0.10) |
| c-Neonicotinoids | 2.80 (4.16 \| 1.44) | 2.66 (3.72 \| 1.60) | 2.69 (3.59 \| 1.79) |
| Organochlorides | 0.98 (2.98 \| -1.02) | 1.1 (3.52 \| -1.32) | 0.33 (2.63 \| -1.97) |

**Table S4:**
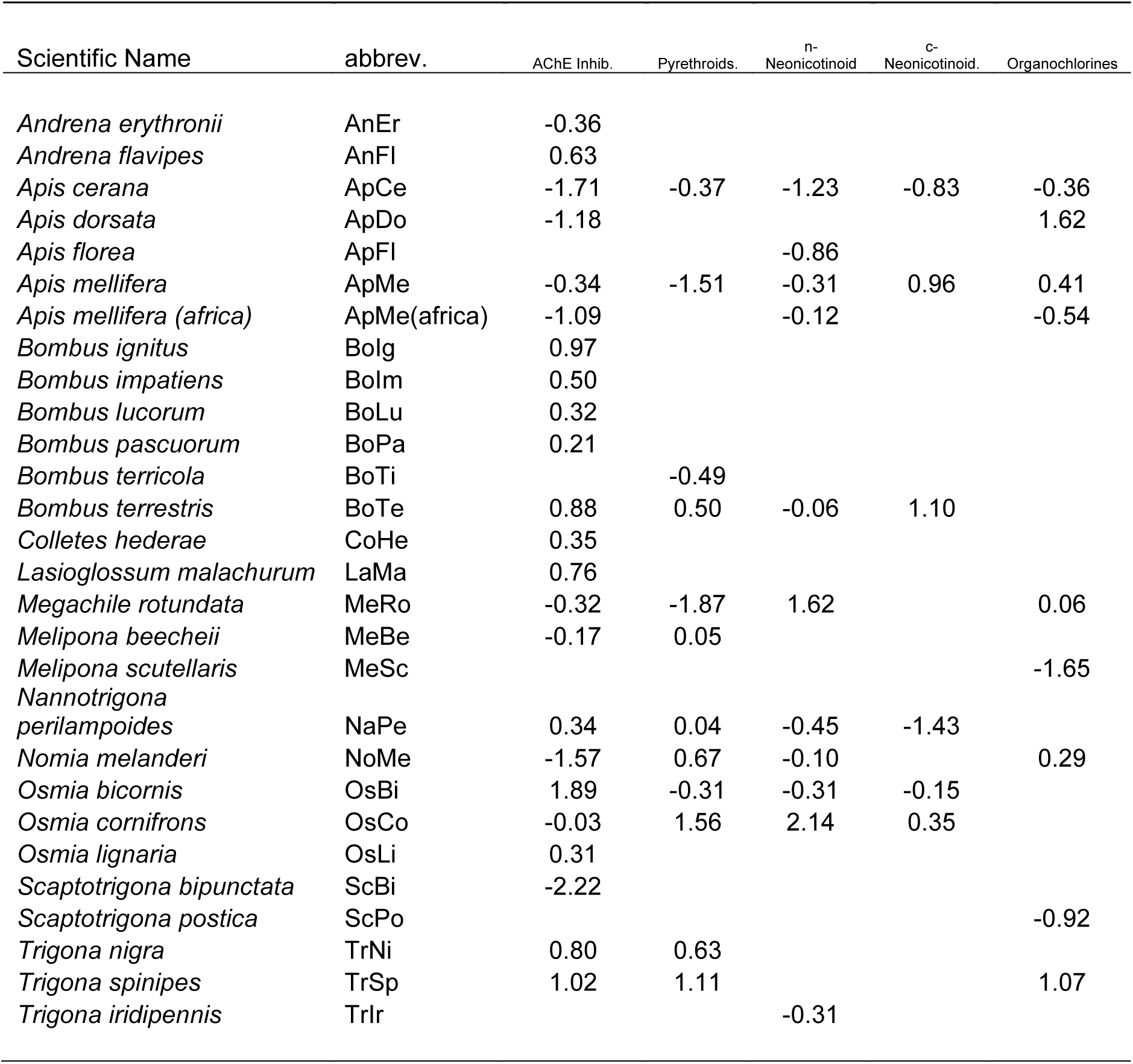
Standardized (z-transformed) relative sensitivity values of bees (per gram) for the five investigated insecticide classes.

## References

1. Ollerton, J., R. Winfree, and S. Tarrant, How many flowering plants are pollinated by animals? Oikos, 2011. 120(3): p. 321–326.

2. Kremen, C., N.M. Williams, and R.W. Thorp, Crop pollination from native bees at risk from agricultural intensification. Proceedings of the National Academy of Sciences, 2002. 99(26): p. 16812–16816.

3. Garibaldi, L.A., et al., Wild pollinators enhance fruit set of crops regardless of honey bee abundance. science, 2013. 339(6127): p. 1608–1611.

4. Biesmeijer, J.C., et al., Parallel declines in pollinators and insect-pollinated plants in Britain and the Netherlands. Science, 2006. 313(5785): p. 351–354.

5. Potts, S.G., et al., Global pollinator declines: trends, impacts and drivers. Trends in ecology & evolution, 2010. 25(6): p. 345–353.

6. Ollerton, J., et al., Extinctions of aculeate pollinators in Britain and the role of large-scale agricultural changes. Science, 2014. 346(6215): p. 1360–1362.

7. Brown, M.J. and R.J. Paxton, The conservation of bees: a global perspective. Apidologie, 2009. 40(3): p. 410–416.

8. Winfree, R., et al., A meta‐analysis of bees’ responses to anthropogenic disturbance. Ecology, 2009. 90(8): p. 2068–2076.

9. Goulson, D., et al., Bee declines driven by combined stress from parasites, pesticides, and lack of flowers. Science, 2015. 347(6229): p. 1255957.

10. EFSA, Scientific Opinion on the science behind the development of a risk assessment of Plant Protection Products on bees (Apis mellifera, Bombus spp. and solitary bees). EFSA Journal, 2012. 10(5): p. 2668.

11. Arena, M. and F. Sgolastra, A meta-analysis comparing the sensitivity of bees to pesticides. Ecotoxicology, 2014. 23(3): p. 324–334.

12. Thompson, H.M. and T. Pamminger, Are honeybees suitable surrogates for use in pesticide risk assessment for non-Apis bees? Pest management science, 2019. 75(10): p. 2549–2557.

13. Lewis, K.A. and J. Tzilivakis, Wild bee toxicity data for pesticide risk assessments. Data, 2019. 4(3): p. 98.

14. Mineau, P., et al., Reference values for comparing the acute toxicity of pesticides to birds. Rev. Environ. Contam. Toxicol, 2001. 170: p. 13–74.

15. Mineau, P., B. Collins, and A. Baril, On the use of scaling factors to improve interspecies extrapolation of acute toxicity in birds. Regulatory Toxicology and Pharmacology, 1996. 24(1): p. 24–29.

16. Davidson, I., J. Parker, and R. Beliles, Biological basis for extrapolation across mammalian species. Regulatory Toxicology and Pharmacology, 1986. 6(3): p. 211–237.

17. Urban, D.H. and N.J. Cook, Hazard evaluation division standard evaluation procedure: Ecological risk assessment. 1986: US Environmental Protection Agency, Office of Pesticide Programs.

18. West, G.B., J.H. Brown, and B.J. Enquist, The fourth dimension of life: fractal geometry and allometric scaling of organisms. science, 1999. 284(5420): p. 1677–1679.

19. West, G.B., W.H. Woodruff, and J.H. Brown, Allometric scaling of metabolic rate from molecules and mitochondria to cells and mammals. Proceedings of the National Academy of Sciences, 2002. 99(suppl 1): p. 2473–2478.

20. West, G.B., J.H. Brown, and B.J. Enquist, A general model for the origin of allometric scaling laws in biology. Science, 1997. 276(5309): p. 122–126.

21. Tomizawa, M. and J.E. Casida, Neonicotinoid insecticide toxicology: mechanisms of selective action. Annu. Rev. Pharmacol. Toxicol., 2005. 45: p. 247–268.

22. Manjon, C., et al., Unravelling the molecular determinants of bee sensitivity to neonicotinoid insecticides. Current Biology, 2018. 28(7): p. 1137–1143. e5.

23. Hansen, T.F. and E.P. Martins, Translating between microevolutionary process and macroevolutionary patterns: the correlation structure of interspecific data. Evolution, 1996. 50(4): p. 1404–1417.

24. Felsenstein, J., Phylogenies and the comparative method. The American Naturalist, 1985. 125(1): p. 1–15.

25. Brady, S.P., et al., Evolutionary toxicology: Toward a unified understanding of life’s response to toxic chemicals. Evolutionary Applications, 2017. 10(8): p. 745–751.

26. Brady, S.P., J.L. Richardson, and B.K. Kunz, Incorporating evolutionary insights to improve ecotoxicology for freshwater species. Evolutionary applications, 2017. 10(8): p. 829–838.

27. Hylton, A., et al., Mixed phylogenetic signal in fish toxicity data across chemical classes. Ecological applications, 2018. 28(3): p. 605–611.

28. Raimondo, S., P. Mineau, and M. Barron, Estimation of chemical toxicity to wildlife species using interspecies correlation models. Environmental science & technology, 2007. 41(16): p. 5888–5894.

29. OECD, Test No. 214: Honeybees, Acute Contact Toxicity Test. 1998.

30. Sparks, T.C. and R. Nauen, IRAC: Mode of action classification and insecticide resistance management. Pesticide biochemistry and physiology, 2015. 121: p. 122–128.

31. Goulson, D., J. Thompson, and A. Croombs, Rapid rise in toxic load for bees revealed by analysis of pesticide use in Great Britain. PeerJ, 2018. 6: p. e5255.

32. DEFRA, Pesticide statistics.

33. DiBartolomeis, M., et al., An assessment of acute insecticide toxicity loading (AITL) of chemical pesticides used on agricultural land in the United States. PloS one, 2019. 14(8).

34. Jeschke, P., et al., Overview of the status and global strategy for neonicotinoids. Journal of agricultural and food chemistry, 2011. 59(7): p. 2897–2908.

35. Hayward, A., et al., The leafcutter bee, Megachile rotundata, is more sensitive to N-cyanoamidine neonicotinoid and butenolide insecticides than other managed bees. Nature ecology & evolution, 2019. 3(11): p. 1521–1524.

36. Beadle, K., et al., Genomic insights into neonicotinoid sensitivity in the solitary bee Osmia bicornis. PLoS genetics, 2019. 15(2): p. e1007903.

37. Del Sarto, M.C.L., et al., Differential insecticide susceptibility of the Neotropical stingless bee Melipona quadrifasciata and the honey bee Apis mellifera. Apidologie, 2014. 45(5): p. 626–636.

38. Motulsky, H., Intuitive biostatistics: a nonmathematical guide to statistical thinking. 2014: Oxford University Press, USA.

39. Michonneau, F., J.W. Brown, and D.J. Winter, rotl: an R package to interact with the Open Tree of Life data. Methods in Ecology and Evolution, 2016. 7(12): p. 1476–1481.

40. Cameron, S.A., H. Hines, and P. Williams, A comprehensive phylogeny of the bumble bees (Bombus). Biological Journal of the Linnean Society, 2007. 91(1): p. 161–188.

41. Danforth, B.N., et al., The impact of molecular data on our understanding of bee phylogeny and evolution. Annual review of Entomology, 2013. 58: p. 57–78.

42. Lo, N., et al., A molecular phylogeny of the genus Apis suggests that the Giant Honey Bee of the Philippines, A. breviligula Maa, and the Plains Honey Bee of southern India, A. indica Fabricius, are valid species. Systematic Entomology, 2010. 35(2): p. 226–233.

43. Rasmussen, C. and S.A. Cameron, Global stingless bee phylogeny supports ancient divergence, vicariance, and long distance dispersal. Biological Journal of the Linnean Society, 2009. 99(1): p. 206–232.

44. Trunz, V., et al., Comprehensive phylogeny, biogeography and new classification of the diverse bee tribe Megachilini: Can we use DNA barcodes in phylogenies of large genera? Molecular Phylogenetics and Evolution, 2016. 103: p. 245–259.

45. Grafen, A., The phylogenetic regression. Philosophical Transactions of the Royal Society of London. B, Biological Sciences, 1989. 326(1233): p. 119–157.

46. Paradis, E., J. Claude, and K. Strimmer, APE: analyses of phylogenetics and evolution in R language. Bioinformatics, 2004. 20(2): p. 289–290.

47. Revell, L.J., phytools: an R package for phylogenetic comparative biology (and other things). Methods in Ecology and Evolution, 2012. 3(2): p. 217–223.

48. Revell, L.J., Phylogenetic signal and linear regression on species data. Methods in Ecology and Evolution, 2010. 1(4): p. 319–329.

49. Orme, D., et al., The caper package: comparative analysis of phylogenetics and evolution in R. R package version, 2013. 5(2): p. 1–36.

50. Team, R.C., R: A language and environment for statistical computing. 2013.

51. Wickham, H., ggplot2: elegant graphics for data analysis. 2016: Springer.

52. Thompson, H., Extrapolation of acute toxicity across bee species. Integrated environmental assessment and management, 2016. 12(4): p. 622–626.

53. Claudianos, C., et al., A deficit of detoxification enzymes: pesticide sensitivity and environmental response in the honeybee. Insect molecular biology, 2006. 15(5): p. 615–636.

54. Michener, C.D., The bees of the world. Vol. 1. 2000: JHU press.

55. Oster, G.F. and E.O. Wilson, Caste and ecology in the social insects. 1978: Princeton University Press.

56. Hughes, W.O., et al., Ancestral monogamy shows kin selection is key to the evolution of eusociality. Science, 2008. 320(5880): p. 1213–1216.

57. Gordon, D.M., The organization of work in social insect colonies. Nature, 1996. 380(6570): p. 121.

58. Dahlgren, L., et al., Comparative toxicity of acaricides to honey bee (Hymenoptera: Apidae) workers and queens. Journal of economic entomology, 2012. 105(6): p. 1895–1902.

59. Free, J., The behaviour of egg-laying workers of bumblebee colonies. The British Journal of Animal Behaviour, 1955. 3(4): p. 147–153.

60. Beig, D., The production of males in queenright colonies of Trigona (Scaptotrigona) postica. Journal of Apicultural Research, 1972. 11(1): p. 33–39.

61. Amsalem, E. and A. Hefetz, The effect of group size on the interplay between dominance and reproduction in Bombus terrestris. PLoS One, 2011. 6(3): p. e18238.

62. OECD, Test No. 246: Bumblebee, Acute Contact Toxicity Test. 2017.

63. OECD, Test No. 247: Bumblebee, Acute Oral Toxicity Test. 2017.

64. Cao, Q., J. Yu, and D. Connell, New allometric scaling relationships and applications for dose and toxicity extrapolation. International journal of toxicology, 2014. 33(6): p. 482–489.

65. Walker, C.H., R. Sibly, and D.B. Peakall, Principles of ecotoxicology. 2016: CRC press.

66. Uhl, P., et al., Is Osmia bicornis an adequate regulatory surrogate? Comparing its acute contact sensitivity to Apis mellifera. PloS one, 2019. 14(8).

67. Uhl, P., et al., Interspecific sensitivity of bees towards dimethoate and implications for environmental risk assessment. Scientific reports, 2016. 6: p. 34439.

68. Moore, D.R., et al., Correcting for Phylogenetic Autocorrelation in Species Sensitivity Distributions. Integrated environmental assessment and management, 2020. 16(1): p. 53–65.

